# Task-based functional connectivity in striato-motor-cortical system in autism: Associations with sex and executive function

**DOI:** 10.64898/2026.03.13.711578

**Authors:** Allison Jack, Abha R. Gupta, GENDAAR Consortium

**Author notes:** **Authors’ contributions.** AJ: Conceptualization, formal analysis, funding acquisition, investigation, methodology, project administration, visualization, writing – original draft; writing – review & editing. ARG: Conceptualization, funding acquisition, investigation, writing – review & editing.

## Abstract

Previously we found that female autistic youth (Aut-F) showed reduced brain response in dorsal striatum (putamen) when viewing human motion, alongside larger rare copy number variants that included genes expressed in early striatal development. Thus, striatal differences may characterize Aut-F, but broader systems-level and behavioral implications of these differences remain unexplored. We conducted secondary data analysis of the sex-balanced cohort (8-17y) in which we first discovered these patterns, in order to: 1) understand how functional connectivity between putamen and frontal targets might vary from the non-autistic population, and differ by sex; and 2) explore which brain connectivity and phenotypic features best predicted executive function. Using psychophysiological interaction analysis (*N*=184), we found that Aut-F youth (*n*=45) showed reduced functional connectivity between left anterior putamen (Pa) and dorsal premotor cortex/pre-supplementary motor area versus matched non-autistic female peers (NAut-F; *n*=45), suggesting reduced engagement of a typical Pa-frontal pathway for attentional regulation. Best subsets regression (*N*=200) indicated that left Pa-left dorsolateral prefrontal functional connectivity explained significant variance in executive functioning across all participants, controlling for neurotype. These results suggest that striatal differences in Aut-F may have adaptive consequences in part due to impacts on connectivity between Pa and frontal regions important for attentional control.

**Lay summary:** We previously found that female autistic people show differences in a part of the brain called the striatum. Some parts of the striatum connect to the frontal lobe of the brain, and may help people control their attention and behavior. We studied how the striatum “talked to” the frontal lobe in autistic girls. We found out that this communication is lower in autistic than non-autistic girls. We also found out that how much striatum “talks to” frontal lobe helps explain differences in how well both autistic and non-autistic youth of both sexes control their attention and behavior.

## Background

Autism diagnosis is higher in male versus female individuals, at rates of ∼3.4 to 1 (Shaw et al., 2025). Autistic people of female gender and/or sex may show different phenotypic characteristics than autistic people of male gender and/or sex (e.g., Cola et al., 2022), and may experience elevated rates of missed or late diagnoses (Aggarwal & Angus, 2015; Bargiela et al., 2016). Given evidence of potential sex- and/or gender-related ascertainment bias (Dworzynski et al., 2012; Russell et al., 2011), identification of objective biomarkers of autism in those assigned female at birth is an important goal for autistic health. In previous work, we observed imaging and genetic evidence that appeared to implicate striatum in the endophenotype and etiology of female autistic youth (Aut-F; Jack et al., 2021). These results suggested that striatum might serve as a promising target for biomarker investigation, but left open several critical questions, including how striatal differences manifested within broader neurocognitive systems. In this short report, we conduct follow-up analyses of these functional magnetic resonance imaging (fMRI) data to probe patterns of striato-motor-cortical system functional connectivity in Aut-F and associations with phenotype.

### Prior work

Previous work with this sample indicated that non-autistic female (assigned sex at birth) youth (NAut-F) engaged executive control regions (e.g., dorsolateral prefrontal cortex [DFC]) to a greater degree than both Aut-F and non-autistic male youth (NAut-M) (Jack et al., 2021). Aut-F demonstrated reduced motor/premotor, dorsal prefrontal, and dorsal striatal (i.e., putamen) fMRI response during social perception, relative to IQ-, head-motion-, and age-matched NAut-F youth. Further, Aut-F, versus autistic male youth (Aut-M), showed significantly greater median size of rare copy number variants (CNVs) containing gene(s) expressed in early development of striatum (Jack et al., 2021), a brain region crucial to acquiring and engaging in motivated behaviors (Grissom et al., 2018). This finding was specific to rare CNVs containing gene(s) expressed in early striatal development, versus those not containing gene(s) expressed in early striatal development. Subsequently, we replicated these genetic findings in the larger SPARK cohort (Jack et al., 2021).

### Striatal system function and sex differences

The striatum is divided into dorsal (caudate; putamen) and ventral (nucleus accumbens) regions. In humans, it contributes to a broad array of functions (linguistic, motoric, affective, social) via integration into associative, limbic, and motor circuits that link regions of striatum to relevant cortical and thalamic regions (Báez-Mendoza & Schultz, 2013; Lapidus et al., 2014). Substantial evidence, particularly in rodents, indicates sex differentiation in striatal neural mechanisms (Meitzen et al., 2018); for example, in dorsal striatum, estradiol mediates increased dopaminergic sensitivity and neuronal firing specifically in animals of female sex (Arnauld et al., 1981). In our fMRI data, sex- and neurotype-related striatal differences were primarily localized in putamen, and previous work in independent (though predominantly male) samples has also documented putaminal differences in autistic versus non-autistic individuals (e.g., Balsters et al., 2018; Di Martino et al., 2011).

### Objectives

We speculated that the neurofunctional and genetic differences we observed in striatal, motor, and frontal sites might be related to each other, indicating variation in the striato-motor-cortical system more broadly.

### Objective 1. Characterize task-specific functional connectivity of the striato-motor-cortical system across sexes and diagnostic groups

We focused on the putamen region of the striatum given this site’s relevance in our previous analyses. We hypothesized that differential putaminal recruitment between Aut-F and NAut-F during social perception might reflect differing organization of functional connectivity, such that this region would be more strongly linked to the executive control network (and associated adaptive and compensatory functions) among NAut-F than Aut-F. Given that, in non-autistic individuals, anterior (Pa) and posterior putamen (Pp) are functionally connected to frontal and motor regions, respectively (Balsters et al., 2018; Morris et al., 2016), we predicted that, for Pa-DFC connectivity: **1A)** NAut >Aut across both sexes and **1B)** NAut-F > NAut-M, reflecting factors related to reduced female likelihood of suprathreshold autistic trait expression. For Pp-M1 connectivity, we expected **1C)** NAut-F > Aut-F, reflecting a disruption to the striato-motor-cortical system underlying the Aut-F functional brain profile previously identified (Jack et al., 2021).

### Objective 2: Explore associations between putamen functional connectivity and executive function

Executive challenges are prominent in autism, where they impact adaptive functioning and may vary by sex (White et al., 2017). Here, we sought to understand how differences in putaminal patterns of functional connectivity to DFC might relate to executive function. DFC is well understood to contribute to executive function through its roles in working memory and attentional control (Jones & Graff-Radford, 2021), and shows functional connectivity to Pa in general population samples (Balsters et al., 2018; Morris et al., 2016). Using an exploratory model building and selection approach, we sought to understand variability in youth executive functioning related to Pa-DFC functional connectivity characteristics.

## Methods

### Participants

We conducted secondary analysis of fMRI data collected from youth (8-17y) in the GENDAAR cohort during passive viewing of coherent (BIO) versus scrambled (SCRAM) point-light displays of human motion. The sample used for group differences analyses was previously described in Jack et al. (2021) as the fMRI sample matched on age (*M*[*SD*]=13.38[2.80]y), full scale IQ, sex, neurotype, and head motion (*N=*184; *n=*45/47 Aut-F/M; *n*=45/47 NAut-F/M; SI Table 1A). For individual differences analyses, all cases from the unmatched fMRI full sample (Jack et al., 2021) with complete BRIEF-GEC data (*M*[*SD*] Aut: 68.63[11.82]; N-Aut: 42.98[6.79]) were used *(N*=200; *n*=44/46 Aut-F/M; *n*=52/58 NAut-F/M; SI Table 1B).

### Data availability

Unprocessed MRI and phenotyping data are available in NIMH Data Archive Collection #2021. Scripts and data needed for replication of study analyses are available at https://osf.io/wekd9/overview?view_only=0ff1800f1d95453d8f98b942d4204679.

### Objective 1

To understand patterns of functional connectivity during social perception, we conducted a psychophysiological interaction (PPI) analysis in FSL using methods described previously (O’Reilly et al., 2012). FMRI quality assurance and preprocessing was identical to that described in Jack et al. (2021). The psychological regressor was the BIO > SCRAM contrast and the physiological regressor was the time course of seeds in the left or right (L, R) Pa or Pp. Seeds were segmented from structural scans using FIRST (Patenaude et al., 2011), with anterior versus posterior division specified per Balsters et al. (2018), using masks kindly provided by the first author (Fig 1A; SI Fig 1). Group differences analyses were conducted within a region of interest (ROI) mask of the dorsal frontal cortex (Brodmann’s Area [BA] 9 & 46d; DFC) from the Sallet Dorsal Frontal Cortex connectivity-based parcellation atlas (Sallet et al., 2013), primary motor (BA 4; M1C) and premotor cortex (BA 6) from the Juelich Histological Atlas (Eickhoff et al., 2005), and insula from the Harvard-Oxford Cortical Atlas (Desikan et al., 2006). Statistical inference used randomise (Winkler et al., 2014) with threshold-free cluster enhancement (Smith & Nichols, 2009), 10,000 permutations, and corrected *p* < .05. Covariates included age, scan site, Social Responsiveness Scale 2^nd^ Edition (Constantino & Gruber, 2012) raw group-mean-centered scores (to account for within-group variability in dimensional autistic traits), and FSIQ.

**Figure 1.**
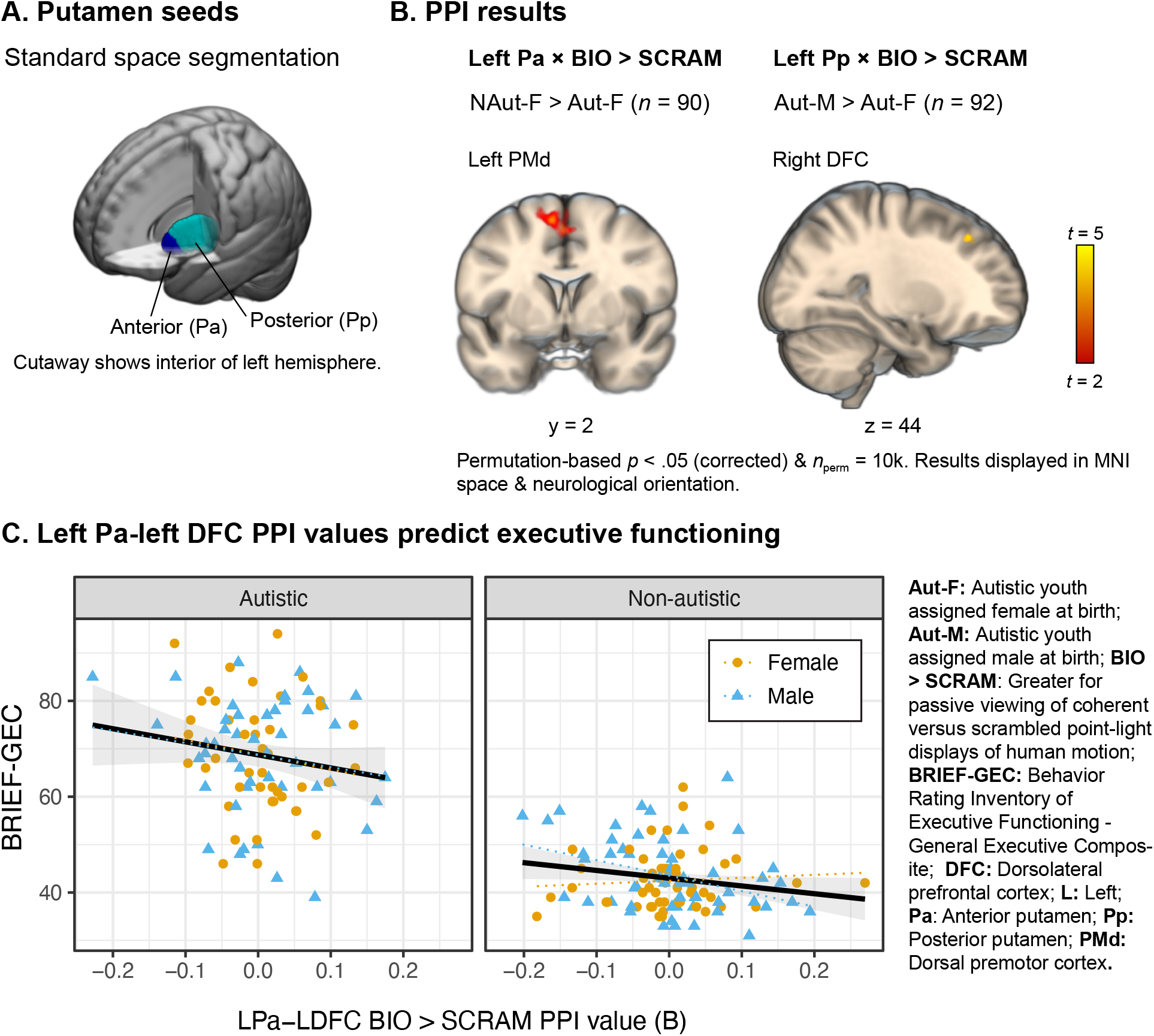
Results of psychophysiological interaction (PPI) group difference (Objective 1) and individual differences (Objective 2) analyses. **A)** A cutaway of the left hemisphere of the Montreal Neurological institute (MNI) standard brain shows the anterior (Pa; dark blue) and posterior putamen (Pp; cyan) masks used for extraction of physiological regressors used in PPI analyses. **B)** Results from PPI group difference analyses, showing sites where the seed showed significantly greater functional connectivity during viewing of coherent vs. scrambled point light displays of human motion (BIO > SCRAM), for one group versus another. Left: non-autistic female youth (NAut-F) showed significantly greater functional connectivity between left Pa and a region of left-to-medial dorsal premotor cortex/pre-supplementary motor area than did autistic female youth (Aut-F). Right: autistic male youth (Aut-M) showed significantly greater functional connectivity between left Pp and right dorsolateral prefrontal cortex (DFC) than did Aut-F. Suprathreshold voxels are colored using a red-yellow gradient using the t-statistic and overlaid on the MNI template. **C)** Individual beta values from the PPI regressor in first level analysis of the interaction between left Pa seed and BIO > SCRAM condition were averaged from across a DFC mask. These values are plotted on the x axis against individual scores on the Behavior Rating Inventory of Executive Function 2nd Ed. – General Executive Composite, where higher scores indicate greater executive deficits. scores are paneled by neurotype, with sex indicated by shape and color. Linear fit lines by group (solid) and sex (dashed) are provided for illustrative purposes.

### Objective 2

We sought the optimal model for predicting individual variability in executive function, as measured via the Behavior Rating Inventory of Executive Function 2^nd^ Edition General Executive Composite (BRIEF-GEC, Gioia et al., 2015). Given the exploratory nature of the objective, we used a best subsets regression (Miller, 2002) approach to feature selection for our model. Features evaluated included fMRI predictors (individual level BIO > SCRAM PPI *b* values with each seed, averaged from within the DFC ROI mask: LPa-LDFC, LPa-RDFC, RPa-LDFC, RPa-RDFC), phenotypic predictors (sex [female = 1]; neurotype [autistic = 1]; age; FSIQ), and interaction terms (all two- and three-way interactions generated from [PPI value]*Sex*Neurotype). A logarithmic transformation was applied to the outcome variable, BRIEF-GEC, to address threats to homoscedasticity. We used leaps v.3.1 in R v.4.2.0 to determine, via exhaustive search, the single best subset of variables for each possible model size (*k*). Repeated cross-validation with 10 folds and 10 repeats (caret v.6.0-92; Kuhn, 2008) was used to estimate root mean squared error (RMSE) for each model, and the model that minimized prediction error was selected as the final model. Standard diagnostics confirmed the final model met linear model assumptions.

## Results

### Objective 1. *During social perception, NAut-F versus Aut-F demonstrate greater functional connectivity between Pa and dorsal premotor cortex*

Partially consistent with our prediction that Aut versus NAut would demonstrate reduced functional connectivity between Pa and targets in frontal cortex, we found that NAut-F versus Aut-F demonstrate greater human-motion selective functional connectivity between left Pa and a region of dorsal premotor cortex sometimes identified as the pre-supplementary motor area (PMd/pre-SMA; Table 1; Fig. 1B, left). However, a similar group difference by neurotype was not observed among male youth.

**Table 1.** Peak voxel coordinates from group difference analyses of PPI between anterior and posterior putamen seeds and regions of frontal, sensory, and motor cortex.

| Seed | Label | Cyt | x | y | z | t | p | k |
| --- | --- | --- | --- | --- | --- | --- | --- | --- |
| **NAut-F > Aut-F** |  |  |  |  |  |  |  |  |
| L Pa | PMd | BA6 | -8 | 2 | 60 | 3.96 | .024 | 158 |
| **Aut-M > Aut-F** |  |  |  |  |  |  |  |  |
| L Pp | DFC/MFG | BA9/46d | 26 | 32 | 44 | 4.83 | .031 | 18 |

**Note.** Group level mixed-effects models controlled for age, FSIQ, scan site, and group-mean-centered Social Responsiveness Scale 2<sup>nd</sup> Edition total raw score. P-values are corrected for multiple comparisons and derived using threshold-free cluster enhancement with 10,000 permutations in FSL's randomise. **Abbreviations.** **Aut-F:** Autistic youth assigned female sex at birth; **Aut-M:** Autistic youth assigned male sex at birth; **BA:** Brodmann's Area; **Cyt:** Cytoarchitectonic; **DFC:** Dorsolateral prefrontal cortex; **k:** Voxel extent; **L:** left; **MFG:** Middle frontal gyrus; **NAut-F:** Non-autistic youth assigned female sex at birth; **Pa:** Anterior putamen; **PMd:** Dorsal premotor cortex; **Pp:** Posterior putamen; **x,y,z:** MNI coordinates.

### During social perception, Aut-M versus Aut-F demonstrate greater functional connectivity between Pp and frontal cortex

Unexpectedly, we found Aut-M to demonstrate greater human-motion-selective functional connectivity than Aut-F between left Pp and right DFC (Table 1; Fig. 1B, right).

### Objective 2. *Social-perception-specific functional connectivity between Pa and DFC predicts executive functioning*

Best subsets regression with repeated cross-validation indicated that a three-variable model predicting BRIEF-GEC scores from neurotype, LPa-LDFC BIO > SCRAM PPI scores, and LPa-LDFC × Sex minimized prediction error (RMSE = 0.16; Adj. R^2^ = 0.67; *F*[3,196] = 136.90; *p* < .001; Table 2; Fig. 1C; see SI Tables 2-3 for predictors selected and model fit parameters for each *k*). The final model indicated that, in addition to the significant variance accounted for by neurotype, greater human-motion-selective functional connectivity between left Pa and left DFC predicted better executive functioning (lower BRIEF-GEC scores).

**Table 2.** Parameters of selected model predicting BRIEF-GEC scores.

|  | **B** | ***t*** | ***p*** |
| --- | --- | --- | --- |
| L Pa-L DFC PPI value | -0.16 | -3.03 | .003 |
| Group | 0.82 | 20.09 | <.001 |
| L Pa-L DFC PPI value × Sex | 0.09 | 1.74 | .083 |

**Note.** L Pa-L DFC PPI value: Average *b* value of the individual level BIO > SCRAM PPI estimate for the left anterior putamen seed, within the dorsal frontal cortex region of interest. Group was coded as Autistic = 1; Non-autistic = 0. Sex was coded as Female = 1; Male = 0.

## Discussion

Our probes of task-based connectivity between putamen and frontal or motor targets revealed three main findings. First, at a group level, NAut-F showed greater social-perception-specific functional connectivity between left Pa and left PMd/pre-SMA than did Aut-F. This region of PMd/pre-SMA has been associated with spatial attention processes (Simon et al., 2002), and is not directly connected to primary motor cortex, but instead to prefrontal sites (Picard & Strick, 2001); therefore, this result is consistent with the previously-reported frontal versus motor distinction between Pa and Pp in neurotypical samples. Greater functional connectivity in this system during social perception suggests that putaminally-driven motivational valuation may upregulate PMd/pre-SMA-mediated spatial attention and gaze control processes during social perception among NAut-F, and that this process may be disrupted or differentially mediated in Aut-F.

Second, Aut-M demonstrated, on average, greater human-motion-selective functional connectivity than Aut-F between left Pp and right DFC. We did not predict this result. However, given that Pp does not typically show functional connectivity to DFC among neurotypical individuals (Balsters et al., 2018; Morris et al., 2016), we interpret this as a potential indicator of greater dysconnectivity in this system in Aut-M versus Aut-F.

Finally, exploratory individual difference analyses indicated that social-perception-specific functional connectivity between left Pa and left DFC significantly predicted variance in general executive abilities as measured by the BRIEF-GEC, even while accounting for executive function differences related to neurotype. In the context of this task, relatively greater L Pa-L DFC connectivity might relate to upregulation of attention during the more “salient” human motion viewing condition. This in turn might be reflective of the participant’s broader attentional control abilities, as reflected in their general executive composite score.

### Limitations and future directions

No information was collected regarding participant sex steroid hormone levels or menstrual cycle phase (where applicable) at the time of fMRI data collection. Unfortunately, this is typical of neuroimaging and biomedical research with menstruating individuals (Schuster & Jansen, 2022). However, animal work indicates that putamen medium spiny neurons are sensitive to estrous cycle phase (Willett et al., 2020). Future work should engage with the challenges of neuroimaging of the menstrual cycle to assess associations between cycle phase and putamen functional connectivity.

### Conclusions & future directions

This project was intended to probe a pattern of results obtained in Jack *et al*. (2021). Future work should pursue this question in an independent sample to determine the replicability of these group and sex differences in putamen functional connectivity. Overall, these findings provide further evidence suggesting putamen-DFC connectivity may be important to adaptive outcomes in autism, and may differentiate Aut-F from NAut-F.

## Acknowledgements

The authors would like to thank Joshua Balsters, PhD, for sharing the anterior and posterior putamen masks used in this study. We would also like to thank all youth who participated in Wave 1 of this study and their families. Additional members of the Wave 1 GENDAAR Consortium MRI team included Kevin A. Pelphrey, PhD; Elizabeth Aylward, PhD; Susan Y. Bookheimer, PhD; Nadine Gaab, PhD; Mirella Dapretto, PhD; and John D. Van Horn, PhD. A full list of Wave 1 GENDAAR team members, with their roles and affiliations, can be found at https://osf.io/mgzny/wiki/0.%20Authors%20%26%20Consortium%20Members/.

---

**Supporting Information Table 1.**
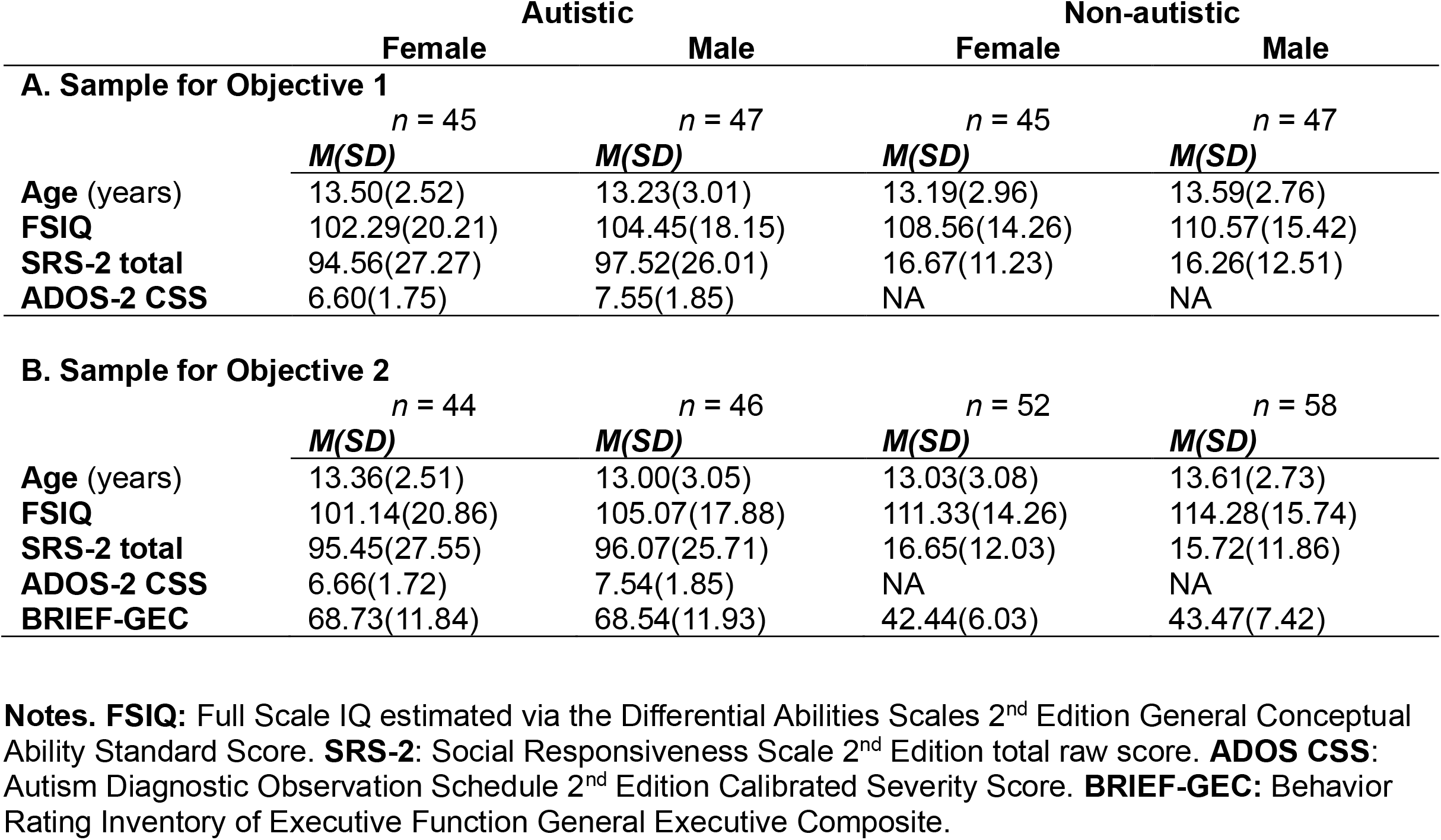
Descriptives for samples used for group differences PPI analysis (Objective 1; A) and best subsets regression (Objective 2; B).

**Supporting Information Table 2.**
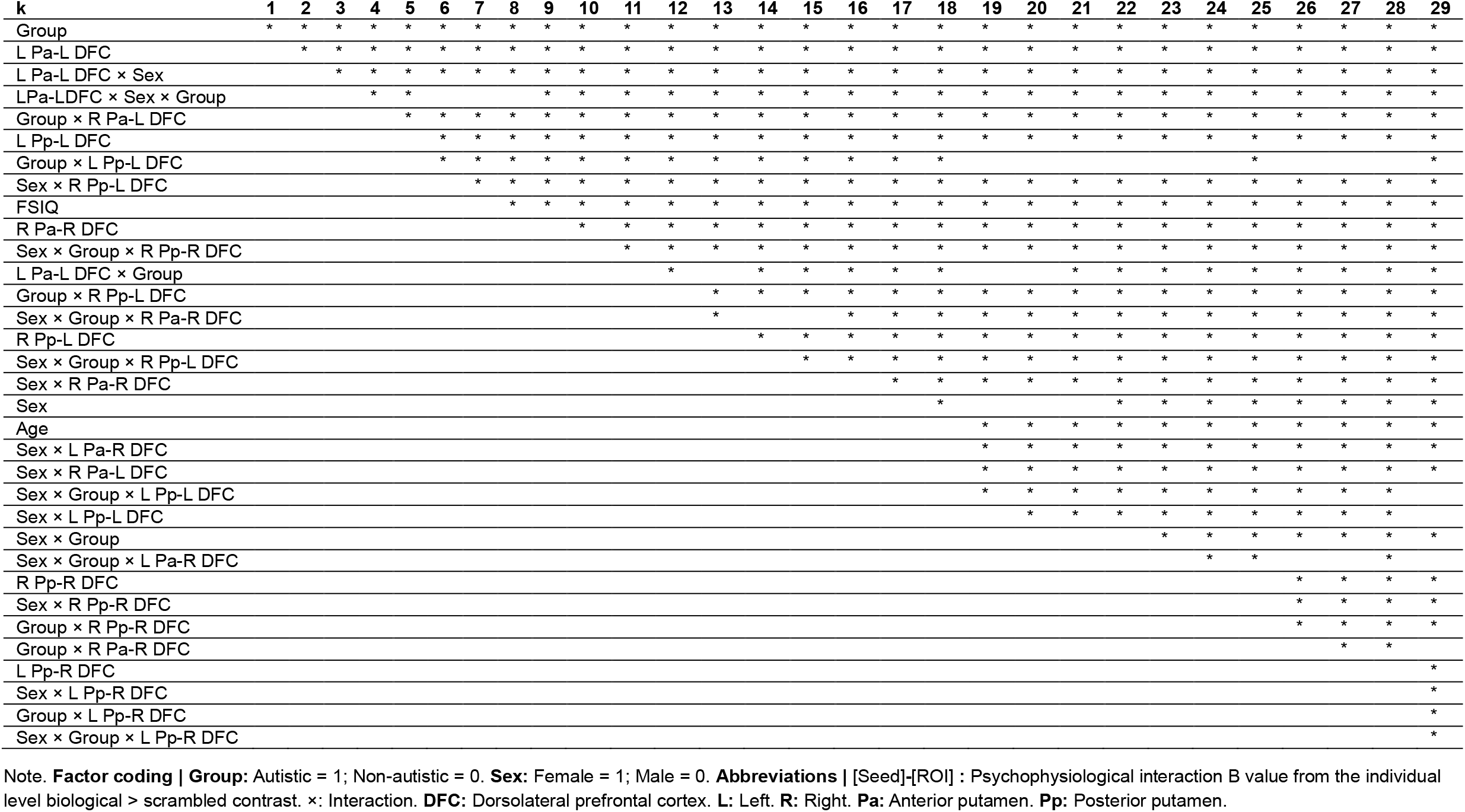
Predictor(s) selected for inclusion at each possible model size (k) by best subsets regression, for prediction of Behavior Rating Inventory of Executive Function – General Executive Composite score.

**Supporting Information Table 3.**
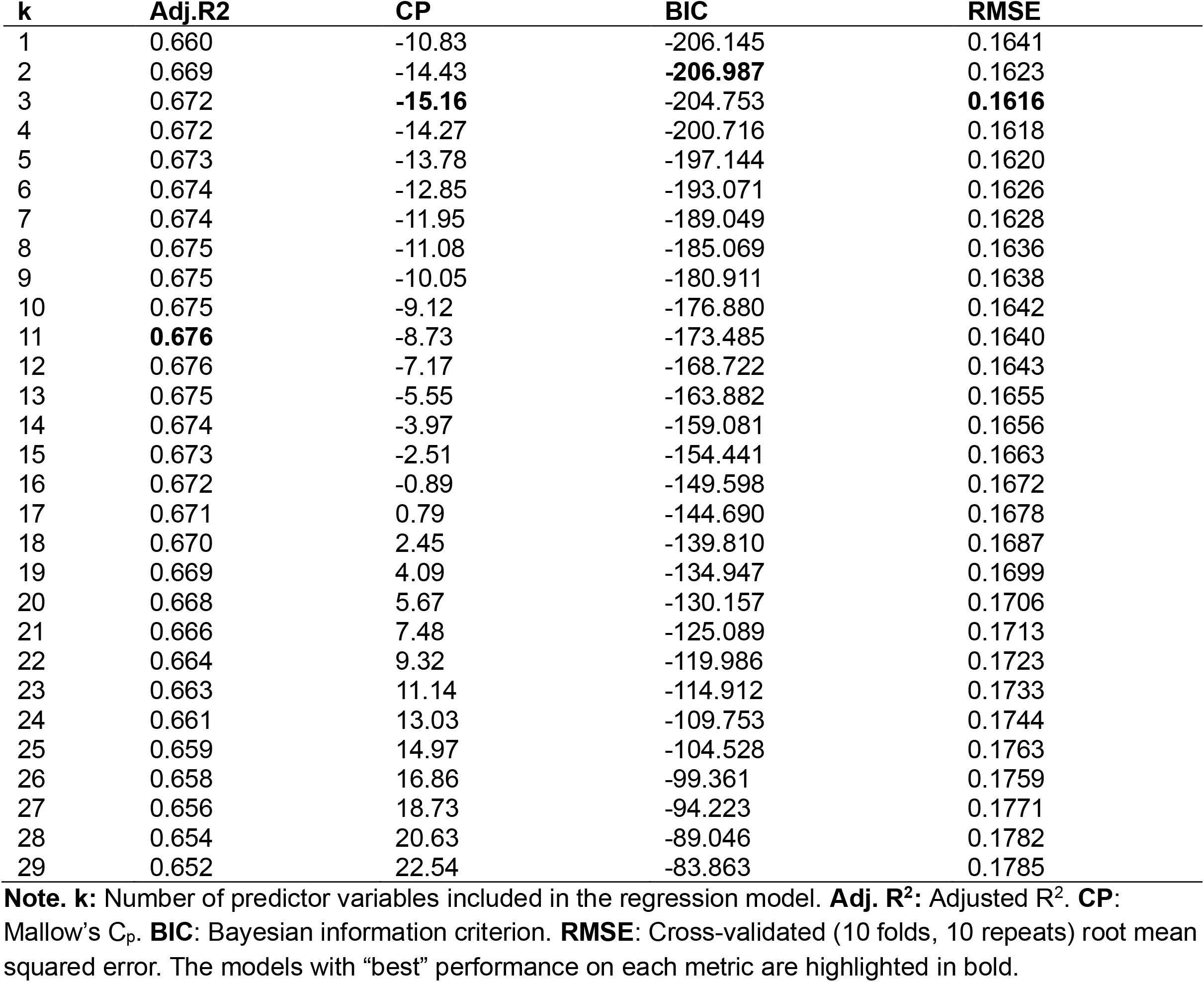
Model performance metrics for each model generated by best subsets regression.

